# Targeting the Cx43 Carboxyl Terminal H2 Domain Preserves Left Ventricular Function Following Ischemia-Reperfusion Injury

**DOI:** 10.1101/668509

**Authors:** Jingbo Jiang, Joseph A. Palatinus, Huamei He, Jegan Iyyathurai, L. Jane Jourdan, Daniel Hoagland, Geert Bultynck, Zhen Wang, Zhiwei Zhang, Kevin Schey, Steven Poelzing, Francis X. McGowan, Robert G. Gourdie

## Abstract

**Background:** αCT1 is a 25 amino acid therapeutic peptide incorporating the Zonula Occludens-1 (ZO-1)-binding domain of connexin43 (Cx43) that is currently in Phase III clinical testing for healing chronic skin wounds. In preclinical studies in mice, we reported that αCT1 reduces arrhythmias and improves ventricular function following cardiac injury, effects that were accompanied by increases in PKCε phosphorylation of Cx43 at serine 368 (pS368). In this study, we undertake a systematic characterization of the molecular mode-of-action of αCT1 in mitigating the effects of ischemia reperfusion injury on ventricular contractile function.

**Methods and Results:** To determine the basis of αCT1-mediated increases in pS368 we undertook tandem mass spectrometry of reactants in an in vitro assay of PKCε phosphorylation, identifying an interaction between negatively charged amino acids in the αCT1 Asp-Asp-Leu-Glu-Iso sequence and positively charged lysines (Lys345, Lys346) in a short α-helical sequence (H2) within the Cx43 CT domain. In silico modeling provided further support of the specificity of this interaction, leading us to conclude that αCT1 has potential to directly interact with both Cx43 and ZO-1. Using surface plasmon resonance, thermal shift and phosphorylation assays, we characterized a series of αCT1 variant peptides, identifying sequences competent to interact with either ZO-1 PDZ2 or the Cx43 CT, but with limited or no ability to bind both polypeptides. Based on this analysis, it was found that only those peptides competent to interact with Cx43, but not ZO-1 alone, resulted in increased pS368 phosphorylation *in vitro* and *in vivo*. Moreover, in a mouse model of global ischemia reperfusion injury we determined that pre-ischemic infusion only with those peptides competent to bind Cx43 preserved left ventricular (LV) contractile function following injury. Interestingly, a short 9 amino acid (MW=1110) Cx43-binding variant of the original 25 amino acid αCT1 sequence demonstrated potent LV-protecting effects when infused either before or after ischemic injury.

**Conclusions:** Interaction of αCT1 with the Cx43 CT, but not ZO-1 PDZ2, explains cardioprotection mediated by this therapeutic peptide. Pharmacophores targeting the Cx43 carboxyl terminus could provide a novel translational approach to preservation of ventricular function following ischemic injury.

## INTRODUCTION

Heart muscle cells are connected together by large numbers of gap junction (GJ) channels ^1, 2^. The main subunit protein of GJs in the mammalian ventricle muscle is Connexin 43 (Cx43 encoded by *GJA1*), which is preferentially localized in intercalated disks – zones of specialized electromechanical interaction between cardiomyocytes ^3, 4^. Following myocardial infarction in patients with ischemic heart disease, Cx43 remodels from its normal distribution in muscle tissue bordering the necrotic injury, redistributing from intercalated disks at cardiomyocyte ends to lateral domains of sarcolemma ^5^. This process of Cx43 lateralized remodeling within the cell membrane is a hallmark of ischemic heart disease in humans and is thought to contribute to the arrhythmia-promoting characteristics of the infarct border zone.

Cx43 phospho-status has emerged as a factor of interest in pathogenic assignments of the protein in the wound healing response of cardiac muscle, and other tissues, including skin ^6^. Pertinent to GJ remodeling in heart disease, Cx43 was observed to be retained at intercalated disks during early ischemia ^7^. This retention occurred in association with increases in phosphorylation at serine 368 (S368) - a consensus Protein Kinase C (PKC) site in the cytoplasmic Carboxyl Terminal (CT) domain of Cx43. Cx43 S368 phosphorylation has also been linked to reduced activity of Cx43-formed channels,^7–9^, including undocked hemichannels ^10^.

Previously, we showed that a peptide mimetic of the Cx43 Carboxyl Terminus (CT), incorporating its postsynaptic density-95/<u>d</u>isks-large/<u>Z</u>O-1 (PDZ)-binding domain reduced Cx43 GJ remodeling in injury border zone tissues following cryo-infarction of the left ventricle in mice ^11^. The decreases in Cx43 remodeling prompted by treatment with this peptide (termed αCT1) were associated with a decreased propensity of the injured hearts to develop inducible arrhythmias ^11^, and sustained improvements in ventricular contractile performance over an 8-week study period ^12^. We further reported that the decreases in Cx43 lateralization observed in hearts treated with αCT1 were correlated with increased phosphorylation of S368 ^11^, in line with results from other workers linking this post-translational modification to reduced GJ remodeling and cardioprotection ^7^.

Initially, we interpreted the induction of increased phosphorylation by αCT1 as a down-stream consequence of the well-characterized property of the peptide to disrupt interactions between Cx43 and its scaffolding protein ZO-1 ^13, 14^. However, in simple biochemical assays involving purified PKC enzyme, and a Cx43 CT substrate, we went on to show that αCT1 promoted S368 phosphorylation *in vitro* in a dose-dependent manner, without recourse to interaction with ZO-1 ^11^. This result raised the prospect that αCT1 mode-of-action could have at least two independent aspects – one involving inhibition of interaction between Cx43 and ZO-1 and the other associated with PKC-mediated changes in Cx43 phospho-status.

The details of αCT1 molecular mechanism is of key translational significance as this therapeutic peptide is presently the subject of testing in the clinic ^15^. In Phase II clinical trials, αCT1 showed efficacy in promoting the healing of two types of chronic, slow healing skin wounds ^16–18^. αCT1 is currently in Phase III testing on more than 500 patients, as a treatment for diabetic foot ulcers (GAIT1 trial) ^19^. In the present study, we provide details of the molecular mechanism of αCT1, showing that the protective effects of αCT1 in ischemic injury to the ventricle is not related to ZO-1 interaction, but is likely associated with binding of the peptide to the Cx43 CT, including the H2 α-helical region - a short stretch of the Cx43 CT adjacent to a serine-rich domain that includes S368.

## MATERIALS AND METHODS

### Animals

Male C57BL/6 mice 3-month old were used. The experimental protocols were approved by Institutional Animal Care and Use Committee of Virginia Polytechnic Institute and State University and conform to the NIH guide for the Care and Usage of Laboratory Animals.

### Reagents: Peptides, cDNA Expression Constructs, and Antibodies

Sequences and a brief description of each Cx43-CT-based peptides used are shown in Table 1. Peptides were synthesized and quality checked for fidelity and purity using High Performance Liquid Chromatography and mass spectometry (LifeTein, Hillsborough, NJ). Biotinylated peptides were designed for surface plasmon resonance experiments. Glutathione-S-Transferase (GST) fusion protein constructs composed of the Cx43-CT (pGEX-6-P2 Cx43 CT amino acids 255-382), ZO-1 PDZ1, PDZ2 and PDZ3 were isolated and purified from isopropy-b-D-thiogalactoside (IPTG)-induced BL21 bacteria using standard procedures, described in our previous publications ^13, 14, 20^. The pGEX6p2-Cx43 CT plasmid was obtained from Prof. Paul L. Sorgen (University of Nebraska Medical Center, USA). Cx43 CT mutant (Cx43 CT-KK/QQ; amino acids Lys345 Lys346 to Gln 345 Gln 346) was developed by site-directed mutagenesis of the pGEX6p2-Cx43 CT plasmid (Agilent technologies, QuikChange II Site-Directed Mutagenesis Kit). The mutation was verified by sequencing. For surface plasmon resonance experiments, the GST was removed using PreScission protease, yielding Cx43 CT protein (wild-type or mutant).

**Table 1.**
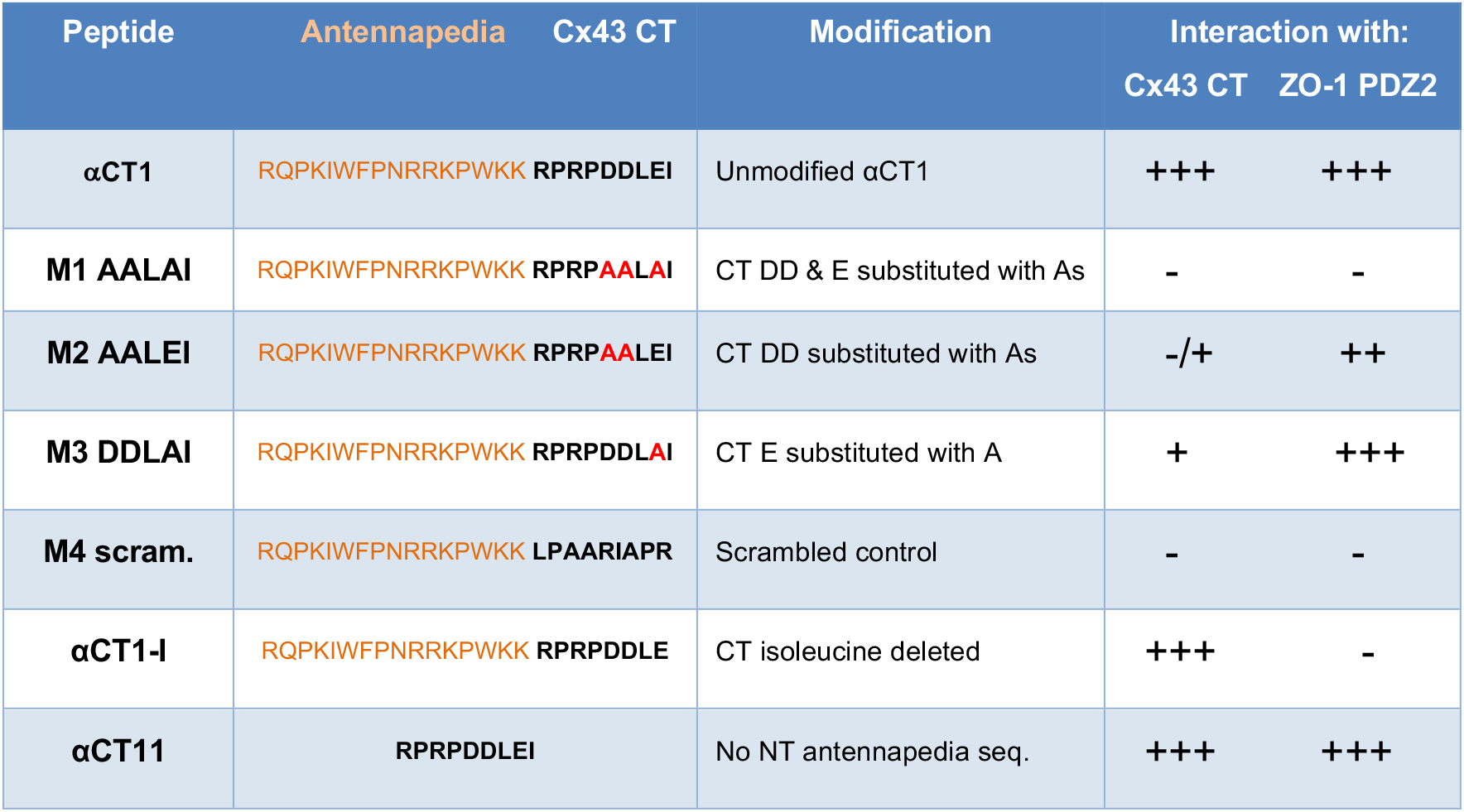
Table of Cx43 mimetic and variant (M) peptide sequences used. Orange letters indicate the antennapedia cell penetration sequence. Red letters indicate alanine substitutions of negatively charged D and E amino acid residues occurring at the CT of wild-type Cx43. The “Interaction with:” column indicates peptide interaction characterization with Cx43 CT or ZO-1, as determined by EDC cross-linking, molecular modeling, SPR and thermal shift assays provided in figures 1 through 4 and supplemental figures 2 and 3.

**Figure 1.**
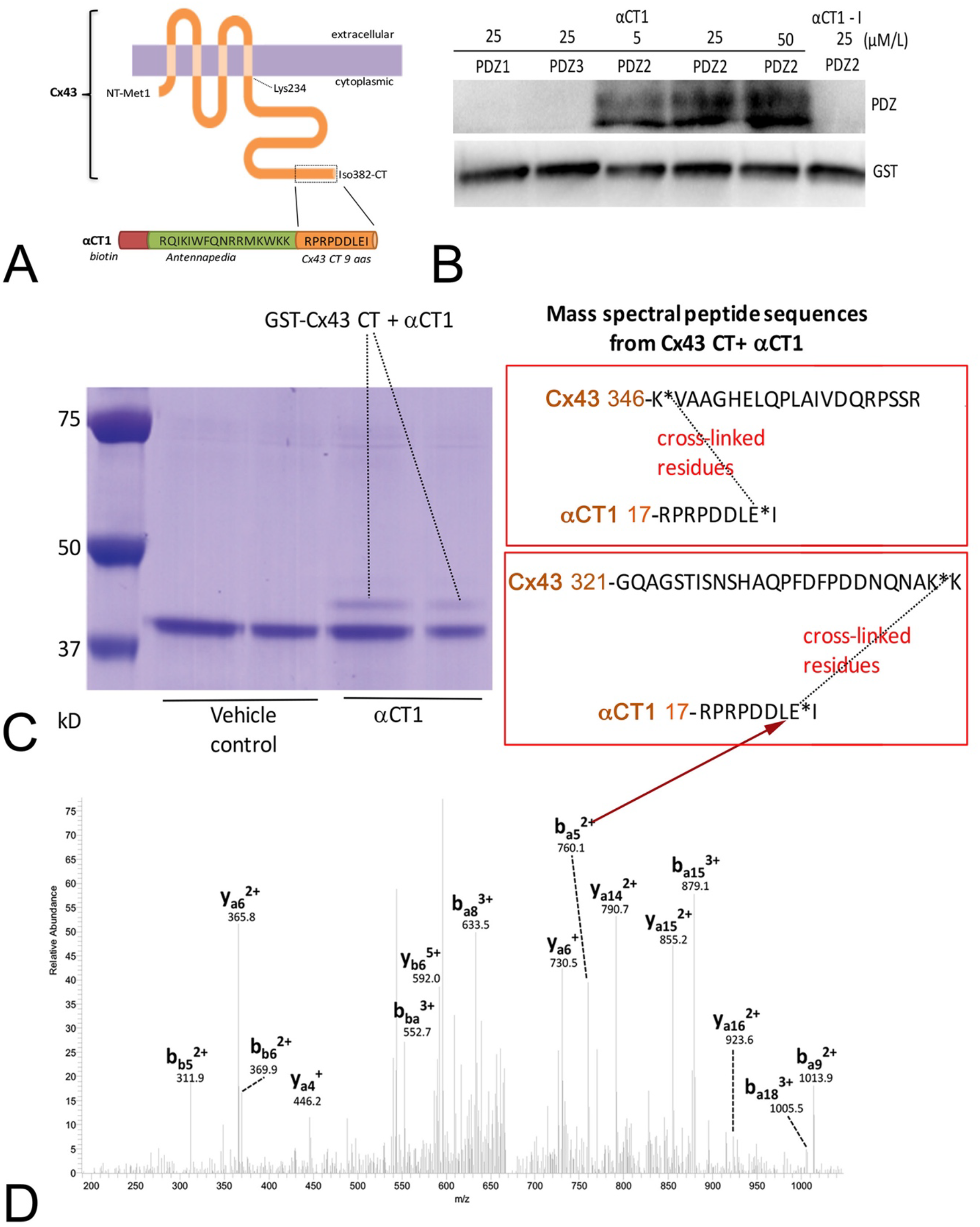
αCT1 interacts with Zonula Occludens-1 (ZO-1) PDZ2 and the Connexin 43 (Cx43) Carboxyl Terminus (CT). **A)** Schematics of full length Cx43 and αCT1 peptide. **B)** αCT1 interaction with ZO-1 PDZ domains as indicated by EDC zero-length cross-linking to GST fusion PDZ1, PDZ2 and PDZ3 polypeptides and neutravidin labeling of biotin-tagged peptide at concentrations of 5, 25 and 50 μM. The deletion of the CT Isoleucine (I) in αCT1-I renders this peptide incompetent to interact with the ZO-1 PDZ2 domain. **C)** Coomassie blue gel of EDC cross-linked products of kinase reaction mixtures containing GST-Cx43 CT and PKC-ε, with (αCT1) and without (Vehicle) αCT1. The fainter band above GST-Cx43 bands (indicated by lines) in the αCT1 lanes were cut from gels and analyzed by Tandem Mass Spectrometry (MS/MS). The boxes to right of gel show Cx43 CT peptides identified by MS/MS as being cross-linked to αCT1. **D)** Tandem mass spectrum of a quintuply charged crosslinked peptide (m/z: 674.1) between Cx43 345-366 (a-chain) and αCT1 peptide through Cx43 K346 and E8 in αCT1 (b-chain). Only the b-and y-sequence specific ions are labeled. Arrow indicates ion (b_a5_^2+^) consistent with cross-linkage between Cx43 CT lysine K346 and the glutamic acid (E) residue of αCT1 at position −1.

#### Antibodies

Phospho-Connexin43 (Ser368) (Cell Signaling, 3511S, Danvers, MA), anti-Cx43 produced in rabbit (Sigma: C6219, St. Louis, MO), anti-GST produced in goat (GE, 27457701, Little Chalfont, UK). NeutrAvidin-HRP (Thermo, 31030, MA).

### Western Blotting

Protein samples from all related experiments (PKC and EDC cross-linking assays and Westerns on heart lysates) were processed in lithium dodecyl sulfate sample loading buffer (Bio-Rad, 1610737 CA), heated at 95°C for 5 minutes. Samples from PKC and cross-linking assays were loaded on 18% Tris-Glycine Stain-Free gels (Bio-Rad, 5678073 CA); samples from heart lysates were loaded on 10% Tris-Glycine Stain -Free gel (Bio-Rad: 5678033 CA), resolved by SDS-PAGE, transferred to PVDF FL membrane on a Turbo Transfer System (Bio-Rad, 1704155 CA). αCT1 eluted from cross-linking reactions was detected on blots against biotin with HRP-NeutrAvidin (ThermoFisher, 31001, MA). Signals were detected by HR-based chemiluminescence (ThermoFisher, 34095, MA) and exposed to ECL Chemidoc (Bio-Rad, 1708280 CA) and digitized using Image Lab software (Bio-Rad, 1709692 CA). Detailed methods have also been previously described in our earlier papers ^11, 13, 14^.

### Surface Plasmon Resonance

Efficacy of the interaction of each αCT1 variant with Cx43 CT or Cx43 CT-KK/QQ was tested using surface plasmon resonance (SPR) as described previously ^20^. In brief, SPR experiments were performed using a Biacore T200 (GE Healthcare). Equal amounts (response units/RU) of biotin-αCT1 variants were immobilized on each flow cell of a streptavidin-coated sensor chip (Biacore Inc) using immobilization buffer (in mM: 10 HEPES, 1 EDTA, 100 NaCl, 0.005% Tween-20) at pH 7.4. Measurements with wild-type (wt) Cx43 CT and mutant Cx43 CT-KK/QQ analytes were done in running buffer (in mM: 10 HEPES, 100 NaCl, pH 7.4) at a flow rate of 30 μl /min. Binding of analytes were verified at different concentrations, in random order (injection volume 120 μl). Interacting proteins were then unbound by injection of 10 μl regeneration buffer (50 mM NaOH and 1 M NaCl) at a flow rate of 10 μl/min. Background levels were obtained from a reference cell containing a biotin-control peptide in which the reversed sequence of the last 9 amino acids of Cx43 was fused to biotin-antennapedia. The RU values obtained with biotin-control peptide were subtracted from the RU values obtained with the different biotin-αCT1 variants (wild-type or mutant) to generate the different response curves.

### PKC-ε Cx43 CT S368 phosphorylation Assay

PKC assay conditions were used to evaluate the PKC**-ε** phosphorylation of Cx43-CT substrate at Ser368 as we have described previously, with modifications ^11^. 400 ng/ml PKC-ε (Life, 37717L, Carlsbad, CA) was pre-diluted in enzyme dilution buffer (10 mM HEPES pH7.4, 0.01% CHAPS and 5 mM DTT) and assayed in 20 mM HEPES pH7.4, 10 mM/L MgCl_2_, 0.1 mM EGTA, 1X lipid mix (200 μg/ml phosphatidylserine (Avanti Polar Lipids 840032C),20 μg/ml Diacylglycerol (Avanti Polar Lipids), 1 mM HEPES pH7.4, 0.03% CHAPS), 500 μM ATP (Sigma, A6419) and 14 μg/ml Cx43-CT substrate. Kinase assay buffer was supplemented with peptides to produce final concentrations of the reaction constituents, as indicated in figure legends. The mixture was incubated at 37°C for 12 minutes and quenched by addition of LDS sample loading buffer (Bio Rad, 1610791). “XT sample buffer” is what was shown on the product label, whereas the component is similar as regular LDS buffer, containing ̴ 5-10% lithium dodecyl sulphate. The reaction was Western blotted for pS368 Cx43 using the Phospho-Connexin43 (Ser368) antibody from Cell Signaling. Proteins were eluted off by stripping buffer (Millipore 2504) and re-probed for total Cx43 using the Sigma anti-rabbit antibody. Percent phosphorylation (% P) was quantified using equation 1 and normalized with control group (PKC+, no peptide added).

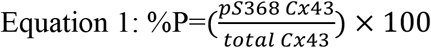

### EDC Cross-Linking Assay

To characterize the interaction between the Cx43 CT substrate and peptides, the in vitro kinase assay was performed as above with modification and the constituents then subjected to a cross-linking reaction. The assay buffer used was 20 mM 3-(N-morpholino) propanesulfonic acid (MOPS), pH 7.2. The Cx43 CT substrate concentration was 30 μg/ml and peptide concentrations varied as indicated in figure legends. All other reagents present in the kinase reaction were maintained as described above. The reaction was allowed to proceed at 37°C for 15 minutes. Afterwards, the carbodiimide crosslinker 1-Ethyl-3-(3-dimethylaminopropyl)carbodiimide HCl (EDC) (Thermo, 22980) was added to each solution for a final concentration of 20 mM. The solution was allowed to cross-link for one hour at room temperature. The reaction was stopped by the addition of 4X LDS loading buffer, boiled for 5 minutes and subsequently separated by PAGE. The resulting gel was stained in Coomassie brilliant blue (Sigma, B0770) for two hours and destained in a solution of 4% methanol 7% acetic acid overnight. Gel bands were subsequently excised for mass spectrometric analysis. For the direct interaction between protein and peptides, PBS, pH 7.5 was used as the coupling buffer. The protein (50 μg/ml) and peptide (25 μM) were allowed to react at room temperature for one hour before EDC was added to the reaction mixture. The reaction was Western blotted for Cx43, GST or NeutroAvidin.

### Tandem Mass Spectrometry

Gel bands corresponding to crosslinked Cx43 and αCT1 were excised and cut into 1 mm square pieces, destained with three consecutive washes with a 50:50 mixture of 50 mM ammonium bicarbonate and acetonitrile for 10 mins. 50 µL of 10 mM DTT was then added to the gel pieces and the gel pieces were incubated at 56°C for one hour. 50 µL of 55 mM iodoacetamide was then added to the sample to alkylate cysteines. The sample was incubated at 25°C in the dark for 45 mins. The gel was then dehydrated with three consecutive washes with a 50:50 mixture of 50 mM ammonium bicarbonate and acetonitrile for 10 min and completely dehydrated with 100% acetonitrile and dried in a speedvac. Gel pieces were rehydrated in 10-15 µL of solution containing 20 ng/µL trypsin (Promega, Madison, WI) in 50 mM ammonium bicarbonate for 15 min. 30 µL of 50 mM ammonium bicarbonate buffer was added to each sample and the samples were incubated at 37°C for 18 hours. Peptides were extracted using 20% ACN/0.1%TFA once, 60%ACN/0.1%TFA twice, and 80%ACN/0.1%TFA once. The extracted samples were pooled and dried in a speedvac and reconstituted in 0.1% formic acid for subsequent LC-MS/MS analysis.

For LC-MS/MS analysis, tryptic peptides were directly separated on a one-dimensional fused silica capillary column (150 mm x 100 μm) packed with Phenomenex Jupiter resin (3 μm mean particle size, 300 Å pore size). One-dimensional liquid chromatography was performed using the following gradient at a flow rate of 0.5 μL/min: 0-10 min: 2% ACN (0.1% formic acid), 10-50 min: 2-35% ACN (0.1% formic acid), 50-60min: 35-90% ACN (0.1%formic acid) balanced with 0.1% formic acid. The eluate was directly infused into an LTQ Velos mass spectrometer (ThermoFisher, San Jose, CA) equipped with a nanoelectrospray source. The instruments were operated in a data dependent mode with the top five most abundant ions in each MS scan selected for fragmentation in the LTQ. Dynamic exclusion (exclude after 2 spectra, release after 30 sec, and exclusion list size of 150) was enabled ^21^.

### Molecular Modeling

Structural information for the Cx43 CT domain truncated at G251 was obtained from the Worldwide Protein Data Bank (DOI:10.2210/pdb1r5s/pdb). The protonated structure of the αCT1 peptide was obtained by truncating the 9 carboxyl terminal amino acids of the Cx43 CT. In order to model the interaction of the Cx43 CT with **α**CT1, the publically available protein-protein docking sever, Zdock (http://zdock.umassmed.edu/help.html) and SWISS-Model were used to model docking of αCT1 with the Cx43 CT in silico in low-energy conformations. Zdock is a Fast Fourier Transform-based protein docking program. Both αCT1 and the Cx43 CT were submitted to the ZDOCK server for possible binding modes in the translational and rotational space. Each pose was evaluated using an energy-based scoring function ^22^.

### Protein Thermal Shift (PTS) Assay

Thermal stability of recombinant GST-PDZ2 or Cx43 CT in the presence or absence of peptides was determined in a 96-well format. Each assay well was composed of 500 μg/mL protein, 25-100 μM of each peptide in PBS buffer, pH7.4. All assays were performed independently six times. Samples were generally prepared in 96-well plates at final volumes of 20 μL. The fluorescent dye SYPRO Orange (5000X concentrate in DMSO, ThermoFisher, S6650) was added to a final concentration of 8X. Reactions were run on QuantStudio 6 Flex Real-Time PCR system (Applied Biosystems, part of Life Technologies Corporation, CA) according to the manufacturer’s recommendations using a melt protocol in 0.05-degree/sec increments from 25°C to 95 °C. The Reporter Dye was “ROX” and quencher Dye and passive reference were selected as “None” for the melt curve according to manufacturer’s instructions. The data were analyzed using Protein Thermal Shift™ Software v1.3 package (Applied Biosystems, CA).

### Ischemia-Reperfusion (I/R) Injury Model and LV Contractility

Male, 3-month-old, body weight 25±5 g C57BL/6 mice were used for this study and obtained from Charles River. Left ventricular (LV) function was measured and myocardial ischemia-reperfusion injury (I/R) was induced as previously described ^23^. Briefly, 15 minutes after the injection of heparin at a dose of 200U/ 10 g body weight, the mouse was anesthetized by inhalation of isoflurane vapor and subjected to cervical dislocation upon the cessation of respiration. Thoracotomy was immediately performed and the heart excised. The heart was arrested in ice-cold Krebs–Henseleit (KH) buffer (in mM: 25 NaHCO_3_, 0.5 EDTA, 5.3 KCl, 1.2 MgSO_4_, 0.5 pyruvate, 118 NaCl, 10 glucose, 2.5 CaCl_2_. The aorta was isolated and cannulated in a Langendorff perfusion system. The heart was then perfused at a constant pressure of 75 mmHg with KH buffer, which was continually bubbled with 5% CO_2_/95% O_2_ at 37°C. Effluent from the Thebesian veins was drained by a thin polyethylene tube (PE-10) pierced through the apex of the LV. A water-filled balloon made of polyvinylchloride film was inserted in the LV and connected to a blood pressure transducer (Harvard Apparatus, 733866, MA). After a 30-minute stabilization period, a balloon volume (BV) generating an LV end-diastolic-pressure (EDP) of 0 mmHg, was determined for the heart. The BV was then increased stepwise up through 1, 2, 5, 8, 10, 12, 15, 18, 20, 25, 30 μl increments of 1-5 μl and contractile performance were recorded for 10 seconds at each step. The indexes of cardiac function were amplified by a Transducer Amplifier Module (Harvard Apparatus, 730065, MA). Data was recorded and analyzed using PowerLab 4/35 (ADInstruments, PL3504, CO) and LabChart V7 (ADInstruments, CO). The BV was then adjusted to set EDP at ∼8-10 mmHg and held constant during the ensuing steps of the protocol. Baseline function (determined by EDP at ∼8-10 mmHg) was recorded for 5 minutes. The perfused beating heart were then treated with freshly prepared peptide stocks (0.2 mM), which were infused using syringe pump (Kent Sci, CT) into the perfusion buffer in a mixing chamber above the heart at 5% of coronary flow rate, to deliver final concentrations of 10-50 μM or equivalent vehicle for 20 minutes. At the end of the peptide infusion period hearts were subjected to global, no-flow normothermic ischemia by turning off the perfusion flow for 20 min, followed by a reperfusion phase for 40 min. BV was retaken through the stepwise sequence of 1-5 μl increments between 1 and 30 μl, with contractile performance again being recorded for 10 seconds at each step. Cardiac LV function was recorded throughout the procedure. In the case of post-ischemic treatment with αCT11 peptide, peptide infusion was begun at the initiation of the reperfusion phase, continued for 20 minutes and then contractile function by BV increments was taken as per the other hearts. A set of hearts were freeze-clamped immediately after peptide infusion for Western blotting. The protocol is illustrated in supplemental figure 1.

### Laser Scanning Confocal Microscopy and fluorescence quantification of peptide perfused hearts

LV samples were Langendorff perfused with vehicle control, αCT1 and αCT11 solutions as described above and as summarized in supplemental figure 1. Immunofluorescent labeling and detection and quantification of biotinylated peptide were performed as previously described ^11, 14, 24^ on 10 µm cryosections of tissue. Samples were co-labeled with a rabbit antibody against either connexin43 (Sigma, C6219, 1:250), Dapi and streptavidin conjugated to AlexaFluor 647 (1:4000; ThermoFisher Scientific). Cx43 primary antibodies were detected by goat anti-rabbit AlexaFluor 488 (1:4000; ThermoFisher) secondary antibodies. Confocal imaging was performed using a TCS SP8 confocal microscope. Quantification of fluorescence intensity levels relative to background were performed using NIH ImageJ software.

### Statistical Analysis

Data were expressed as a mean ± SE unless otherwise noted. Differences among treatments were compared by one-way, two-way or repeated measures ANOVA, followed by post hoc or Mann-Whitney tests, as appropriate. Probability values *p* < 0.05 were considered significantly different. No strong evidence of divergence (p>0.05) from normality was found. Data analysis was performed using GraphPad7 (GraphPad Software, LaJolla, CA).

## RESULTS

### αCT1 interacts with the Cx43 Carboxyl Terminus H2 domain

The 25mer Cx43 mimetic peptide αCT1 incorporates a 16-amino acid N-terminal (NT) antennapedia (Antp) sequence followed by the carboxyl terminal (CT)-most 9 amino acids of Cx43: Arg-Pro-Arg-Pro-Asp-Asp-Leu-Glu-Iso or RPRPDDLEI (Fig. 1A and Table 1). The last four amino acids of this sequence (DLEI) comprise a class II PDZ-binding motif, which has been shown to mediate a specific interaction with the second of the three PDZ (PDZ2) domains of ZO-1 ^14, 25, 26^. We have previously reported on binding of αCT1 with ZO-1, and the selectivity of this interaction for the ZO-1 PDZ2 domain over that of ZO-1 PDZ1 and PDZ3 ^14^. This selectivity of αCT1 for ZO-1 PDZ2 is illustrated in Fig. 1B. Consistent with reports by others ^27^, deletion of the CT isoleucine of the DLEI binding motif (e.g., as in the αCT1-I peptide, Table 1) abrogates interaction with ZO-1 PDZ2 (Fig. 1B). We also have previously shown that αCT1 upregulates a PKCε-mediated phosphorylation of Cx43 at serine 368 (S368) along its primary sequence ^11^. This induction of S368 phosphorylation (pS368) by αCT1 was observed both *in vivo* in a left ventricular (LV) injury model and in a biochemical assay of PKCε activity *in vitro* ^11^.

To identify the molecular determinants of αCT1-induced upregulation of S368 phosphorylation, the zero-length cross-linker 1-ethyl-3-(-3-dimethylaminopropyl) carbodiimide hydrochloride (EDC) was introduced into the in vitro PKCε phosphorylation assay. Zero-length cross-linking covalently bonds directly interacting proteins, enabling identification of partnering proteins in the reaction mixture. The components of the reaction mixture were then separated by SDS-PAGE and tandem mass spectrometry (MS/MS) was performed on the isolated polypeptides (Figs. 1C and 1D). While no evidence for interaction between PKCε and αCT1 was observed, the analysis revealed that a band running just above GST-Cx43 CT corresponded to a covalently linked complex between the Cx43 CT substrate and αCT1 (Fig. 1C). Moreover, it was determined that a negatively charged glutamic acid (E381) within the PDZ-binding domain of αCT1, and a pair of aspartic acids (D378, D379), were involved in bonding with a pair positively charged lysines (K) at positions K345 and K346 of the Cx43 CT (Figs. 1C and 1D). The site specificity of this interaction was further confirmed by streptavidin labeling of cross-linked products from kinase reaction mixtures containing Cx43 CT with αCT1 (Supplemental Fig. 2 left hand blots) or a scrambled peptide (M4) unable to bind Cx43 CT (Supplemental Fig. 2,, middle blots), as well as in reaction mixtures containing a mutated Cx43 CT (GST-Cx43 CT QQ/KK) substrate (Supplemental Fig. 2 right hand blots), in which the pair of positively charged lysine residues at K345 and K346 were substituted with neutral glutamines (Q). While no evidence of cross-linking between the scrambled peptide and Cx43 CT, or between αCT1 and the Cx43 CT QQ/KK mutant substrate was found, αCT1 was covalently linked by EDC to Cx43 CT in a concentration-dependent manner.

### αCT1 interacts with the Cx43 CT H2 domain

Structural studies by Sorgen and co-workers have shown that K345 and K346 fall within a short α-helical sequence along the Cx43 CT called H2 (for Helix 2) ^28, 29^. Figure 2A provides a schematic of the secondary structure of the Cx43 CT showing the location of H2 (from ^30^), together with a second nearby stretch of the α-helical sequence (H1). To model αCT1:Cx43 CT H2 binding in silico, we submitted the interacting complex to the zDOCK protein modeling server ^22^, initially fixing the interaction between the glutamic acid (E) at position −1 of αCT1 (i.e., E381 in full length Cx43) and the K346 residue of Cx43 – as predicted by the MS/MS data (i.e., Fig. 1C). The interaction pose shown in Figure 2B represents that based on the lowest energy minimization score from over 1800 possible variants of the complex. Using Schrodinger molecular modeling software, and without specifying the initial H2 K346 e381 of αCT1 bonding constraint, we confirmed that αCT1 could be optimally configured in an anti-parallel orientation with its available side-chains arrayed along the H2 sequence (Figs. 2C, D). As indicated by MS/MS, a salt bridge was predicted by this *in silico* analysis to form between the αCT1 glutamic acid (E) residue and Cx43 K346. The modeled interaction further anticipates hydrogen bonding between the side chains of four amino acids arrayed along αCT1 (**R**P**R**PD**D**LE**I** – amino acids involved in hydrogen-bonds are bolded) and four amino acids between Q340 and E360 of the H2 sequence (Fig. 2D).

**Figure 2.**
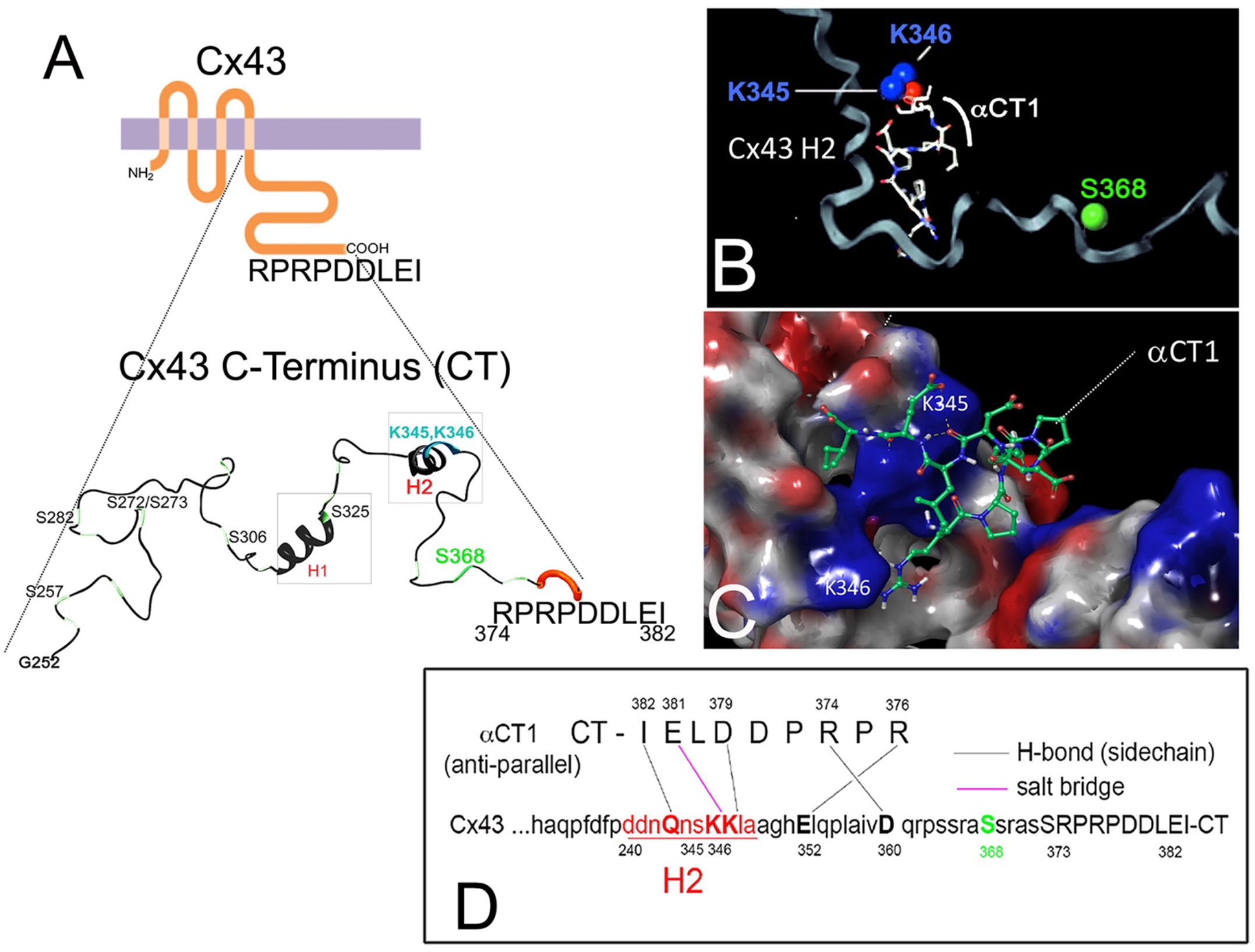
Molecular modeling of the αCT1 and Cx43 CT complex. **A)** Schematics of Cx43 and the secondary structure of Cx43 CT from amino acid residues Glycine252 (G252) through to Isoleucine 382 (I382). The depiction of secondary structure in 2A has been modified from a diagram originally provided by Sosinsky and co-workers ^30^. **B)** ZDOCK and **C)** Schrodinger molecular modeling software analysis of the structure of a proposed αCT1-Cx43 CT complex. The protonated structure of αCT1 peptide and Cx43 CT (PDB:1r5s), constrained by a salt-bridge interaction between K346 in the Cx43 CT and the glutamic acid (E) at position −1 of αCT1. The αCT1-Cx43 interaction shown represents that based on the lowest energy minimization score determined in the model. **D)** Schrodinger molecular modeling software, a 2D map of αCT1-Cx43 CT in anti-parallel orientation showing location of amino acids predicted to bond to each other and the type of bond that is predicted to occur.

### Substitution of negatively charged amino acids in αCT1 results in loss of Cx43 CT binding

To further probe the αCT1 complex with Cx43 CT H2 region, and its consequence for phosphorylation of S368, three variant peptides based on αCT1 were prepared. In these peptides, negatively charged E and D amino acids in the RPRPDDLEI sequence of αCT1 (i.e., those indicated by MS/MS to be likely involved in Cx43 CT interaction) were substituted by neutral alanines. These αCT1 variant peptides had the sequences RPRPAALAI, RPRPAALEI, and RPRPDDLAI and are referred to as M1 AALAI, M2 AALEI and M3 DDLAI respectively. First, surface plasmon resonance (SPR) was used to analyze interactions of biotinylated versions of αCT1 and the αCT1 variants peptides, immobilized to streptavidin-coated sensor chips, with the Cx43 CT and Cx43 CT-KK/QQ proteins as analytes (Figs. 3A-F, Supplemental Fig. 3A, B). The concentration of the analyte was varied between 0.5 and 15 μM. A concentration-dependent increase in Response Units was observed for Cx43 CT binding to biotin-αCT1 (Fig. 3A). M1 AALAI showed loss of Cx43 CT binding competence, consistent with having all negatively charged amino acids substituted with alanine (Fig. 3C). Substitution of D378/D379 (M2 AALEI) or E381 (M3 DDLAI) residues by alanines also abrogated peptide interaction with Cx43 CT (Figs. 3E, F, Supplemental Fig. 3A, B). In complementary observations, SPR confirmed that the Cx43 CT KK/QQ mutant polypeptide was unable mediate interactions with αCT1, M1 AALAI, M2 AALEI or M3 DDLAI (Figs. 3B, D, and F), consistent with the pair of lysines at K345 and K346 in H2 being necessary for interaction between Cx43 and αCT1.

**Figure 3.**
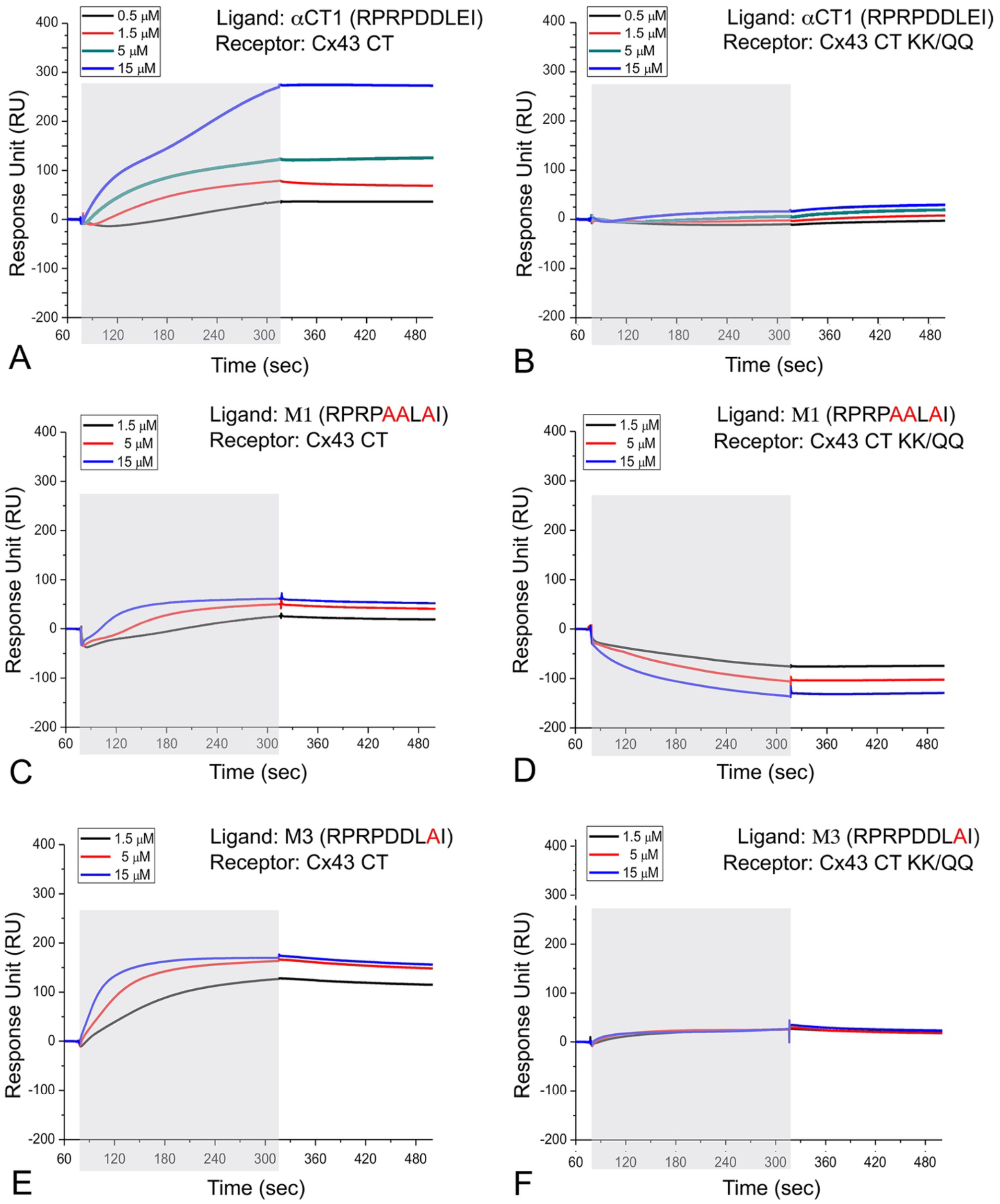
αCT1 variants with alanine substitutions of negatively charged amino acids show abrogated ability to bind Cx43 CT (3A-F). SPR was used to analyze interactions of biotin-αCT1 and biotin-αCT1 variant peptides, immobilized to streptavidin-coated chips, with the Cx43 CT (Cx43-CT: amino acids 255 to 382) and Cx43 CT-KK/QQ as analytes, respectively. The mean of three runs is plotted for each analyte concentration. The exposure of the sensor chip to the specific analyte is indicated by the gray area. Sensorgrams obtained for: **A)** Cx43 CT and biotin-αCT1. **B)** Cx43 CT-KK/QQ and biotin-αCT1. **C)** Cx43 CT and biotin-M1 AALAI. **D)** Cx43 CT-KK/QQ and biotin-M1 AALAI. **E)** Cx43 CT and biotin-M3 DDLAI. **F)** Cx43 CT-KK/QQ and biotin-M3 DDLAI.

### Substitution of negatively charged amino acids in αCT1 fully and partially abrogate interaction with Cx43 CT and ZO-1 PDZ2 respectively

To further characterize the Cx43-binding characteristics of αCT1 and the αCT1 variants, thermal shift assays of peptide:protein interactions were performed (Figure 4). This assay provides quantitative data on the effect of interaction on protein structure – with significantly increased or decreased thermal stability in response to temperature cycles in a qPCR machine being diagnostic of potential interaction. For example, in line with the known stabilizing effect of the last 10 amino acids of Cx43 CT on ZO-1 PDZ2 ^31^, αCT1 concentrations of 25, 50 and 100 μM increased the melt temperature (i.e., thermal stability) of PDZ2 in a dose-dependent manner (Fig. 4A). Thermal shift assays indicated that the peptides from Table 1 fell into two classes with respect to Cx43 CT interaction – those that provided evidence of interaction with Cx43 CT and those that were Cx43 CT interaction incompetent (Fig. 4B). Consistent with the SPR results, M1 AALAI, M2 AALEI and M3 DDLAI showed no propensity to alter Cx43 CT thermal stability, demonstrating no significant variance from Cx43 CT alone or Cx43 CT in the presence of the scrambled control peptide M4. By contrast, αCT1, αCT1-I and short variant of αCT1 comprising the Cx43 CT 9mer sequence RPRPDDLEI (αCT11), all caused highly significant decreases in melt temperature, in line with interaction of these peptides disrupting protein thermal stability via binding to the Cx43 CT.

**Figure 4.**
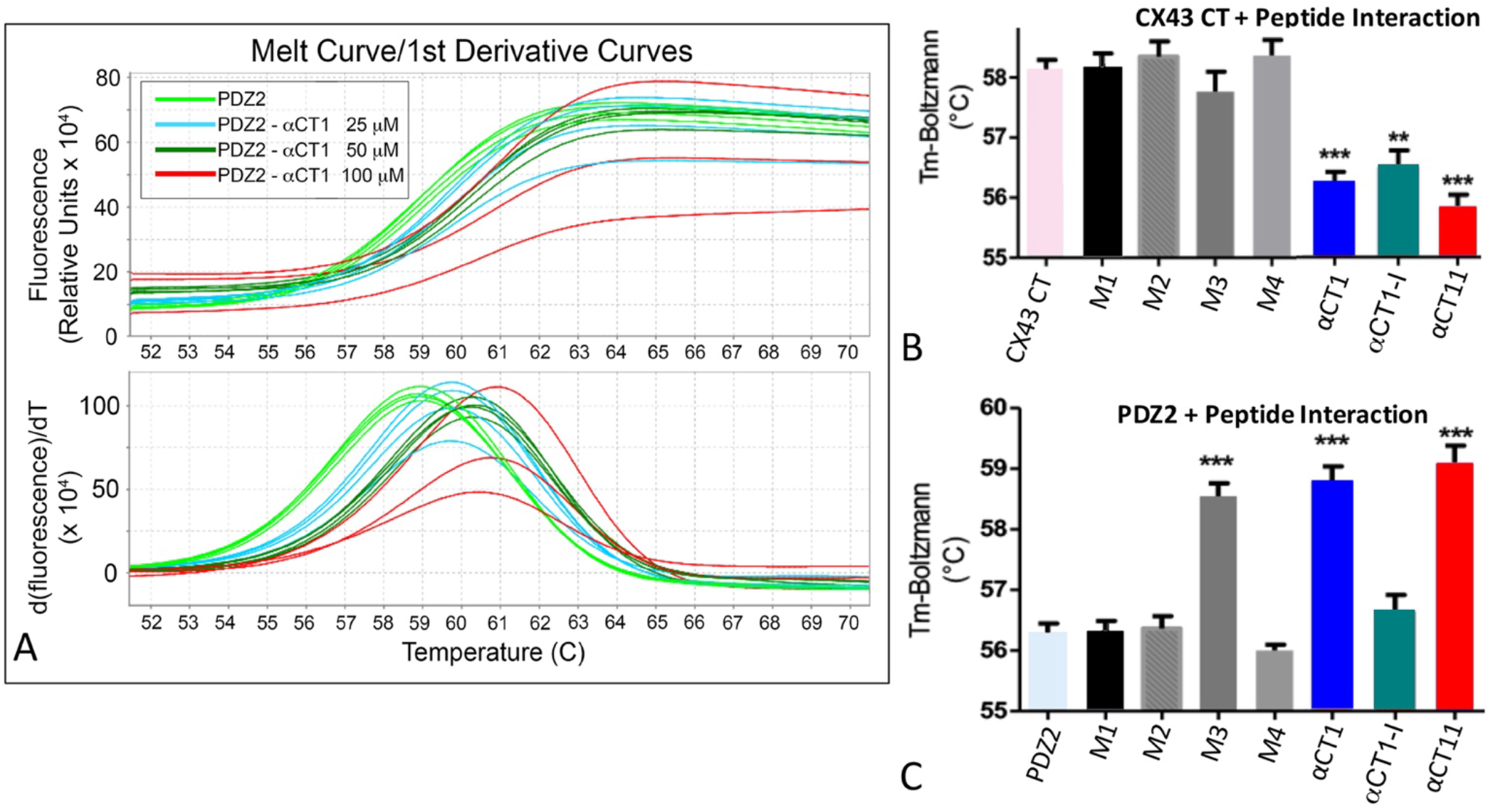
αCT1 interaction stabilizes PDZ2 and destabilizes Cx43 CT secondary structure. **A)** Melt curves (top) and first derivative of melt curves (bottom) for ZO-1 PDZ2 at 500 μg/mL in combination αCT1 at concentrations of 25, 50 and 100 μM. **B)** Temperature maxima (Tm) from Boltzman curves from left-to-right of Cx43 CT (Cx43-CT: amino acids 255 to 382) alone, Cx43 CT in combination with αCT1, and the αCT1 variants including: M1 AALAI, M2 AALEI, M3 DDLAI, M4 scrambled, αCT-I and αCT11. αCT1, αCT1-I and αCT11 show similar abilities to destabilize (i.e., significantly decrease the Tm of) Cx43 CT. **p<0.01, *** p<0.002, N=6. **C)** Temperature maxima (Tm) from Boltzman curves from left-to-right of PDZ2 alone, and PDZ2 in combination with αCT1 and αCT1variants including αCT1 variants including: M1 AALAI, M2 AALEI, M3 DDLAI, M4 scrambled, αCT-I and αCT11. M3 DDLAI, αCT1, and αCT11 show similar abilities to stabilize (i.e., significantly increase the Tm of) PDZ2. **p<0.01, ***p<0.002, N=6

We also examined the effects of the αCT1 variants on thermal stability of ZO-1 PD2. Unlike in presence of the parent peptide αCT1 (Fig. 4A, C), M1 AALAI and M2 AALEI did not alter the melt temperature of PDZ2 (Fig. 4C), not differing significantly from PDZ2 alone, or PDZ2 in the presence of either scrambled peptide (M4) or αCT1-I -the two PDZ2 interaction incompetent peptides (Fig. 1). The results were consistent with M1 AALAI or M2 AALEI having no, or limited, propensity to interact with the ZO-1 domain. However, M3 DDLAI, the most conservative substitution variant, showed evidence of significant interaction with its ZO-1-binding domain, with its effects on the thermal stability of PDZ2 being similar in this assay to those of αCT1 (Figure 4C). Thus, although M3 DDLAI had no or limited competence to interact with Cx43 CT, this peptide did show evidence of ZO-1 PDZ2-binding activity not significantly different from unmodified αCT1.

### Substitution of negatively charged amino acids in αCT1 abrogates induction of S368 phosphorylation

Next, we examined how the mutant peptides performed in the PKC-ε kinase assay. Unlike αCT1, neither M1 AALAI, M2 AALEI nor M3 DDLAI increased Cx43 S368 phosphorylation above levels detected in the absence of peptide (PKCε+plus lanes of Fig. 5A), or in the presence of scrambled control peptide (M4, Fig. 5A, B). Quantification of blots indicated that the ability of unmodified αCT1 to induce S368 phosphorylation was ~ 3-fold greater than that of either the PKCε+plus control reaction (p<0.001) or reactions including M1 AALAI or M4 peptides (Fig. 5C). It was further determined that a 9 amino acid peptide comprising only RPRPDDLEI (i.e., αCT11 = αCT1 with its 16 amino acid NT antennapedia sequence truncated) robustly upregulated pS368 levels over control (vs PKCε+plus control <0.001) (Fig. 5C). αCT1-I, the ZO-1-binding-deficient peptide with CT isoleucine truncated, also prompted a significant increase in PKCε-mediated phosphorylation of Cx43 CT (p<0.05 vs. PKCε+plus control). In sum, the results indicated that only those αCT1-based peptides competent to interact Cx43 CT (i.e., αCT1, αCT11 and αCT1-I), but not those unable to (i.e., M1 AALAI, M2 AALEI, M3 DDLAI and M4), increased pS368 above control levels. Also, given that M3 DDLAI is unable to induce pS368 increase, but does retain PDZ2 interaction ability, the data suggested that ZO-1-binding activity is dispensable for this phosphorylation.

**Figure 5.**
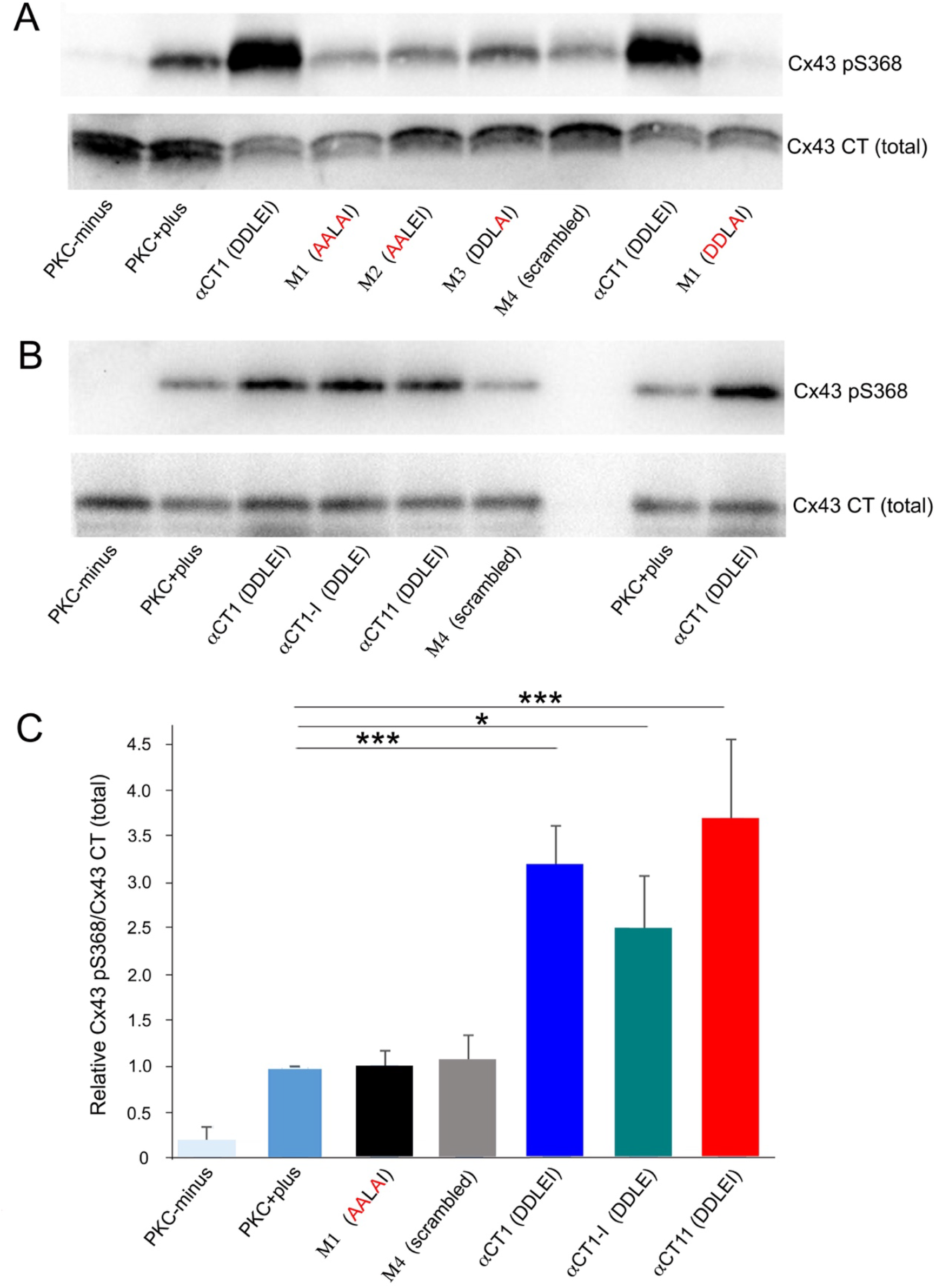
Cx43 mimetic peptides that retain Cx43-binding capability are able to induce phosphorylation of Cx43-CT at serine 368 (S368). **A)** Blots of Cx43-pS368 (top) and total Cx43 (bottom) in kinase reactions mixtures including no-kinase controls with substrate (Cx43-CT: amino acids 255 to 382), but no PKC-ε (PKC-minus); Cx43-CT substrate with PKC-ε (PKC-plus); and mixtures containing PKC-ε, Cx43 CT, and biotin-tagged αCT1, biotin-tagged αCT1 mutant peptides with alanine substitutions (M1 AALAI, M2 AALEI, M3 DDLAI) and biotin-tagged M4 scrambled. Peptides are at 20 μM. **B)** Blots of Cx43-pS368 (top) and total Cx43 (bottom) in kinase reactions mixtures including no-kinase controls with Cx43 CT substrate, but no PKC-ε (PKC-minus); Cx43-CT substrate with PKC-ε (PKC-plus); and mixtures containing PKC-ε, Cx43 CT, and biotin-αCT1, biotin-αCT1-I or biotin-αCT11 (RPRPDDLEI with no antennapedia sequence at peptide NT) and biotin-M4 scrambled peptide. Peptides are at 20 μM. **C)** Chart showing that the ability of unmodified αCT1 and the Cx43 CT interaction-competent peptides biotin-αCT1-I or biotin-αCT11 to induce S368 phosphorylation was 3-5 fold greater than that of non-Cx43 CT interacting peptides. * p<0.05, ** p<0.01, *** p<0.002, N=5 αCT1 and M4, other peptides N=3.

### Only peptides interacting with Cx43 CT protect hearts from ischemic injury

The biochemical characterizations indicated that αCT1 is capable of two distinct protein-protein interactions - one with ZO-1 PDZ2 and the other with the Cx43 H2 region. This raised the question as to whether or not the previously characterized effects of αCT1 in cardiac injury models ^11, 12^, or indeed its wound healing effects at large ^16–18, 24^, could be accounted for by one or another of these protein-protein interactions. The series of αCT1-based variant peptides generated for the present study provided an opportunity to address this question. While αCT1-I is not competent to interact with ZO-1 PDZ2, this αCT1 variant does bind the Cx43 CT and upregulate S368 phosphorylation. Conversely, while M3 DDLAI showed no ability to bind Cx43 CT, or increase pS368, this peptide retained affinity for the ZO-1 PDZ2 domain. Finally, M1 AALAI showed no evidence of interaction with either PDZ2 or Cx43 CT, and demonstrated no ability to increase pS368 in the in vitro assay. We thus used the variant peptides, together with unmodified αCT1 in mouse hearts subjected to an ischemia-reperfusion (I/R) protocol to systematically assess which aspect of mode-of-action (i.e., peptide interaction with ZO-1 vs. Cx43) accounted for modulation of the I/R injury response by Cx43 CT mimetic peptides.

The protocol and experimental design for the cardiac I/R injury model is illustrated in supplementary Figure 1. In summary, the protocol involved a 20-minute period of no flow ischemia period followed by 40 minutes of reperfusion. For treatment, peptides were infused into hearts over a 20-minute period just prior to the ischemic episode. Representative pressure traces from a vehicle control and αCT1-treated hearts are shown in Figures 6A and B, from which it can be qualitatively appreciated that pre-ischemic infusion of αCT1 results in preservation of LV contractile function upon reperfusion relative to vehicle control.

**Figure 6.**
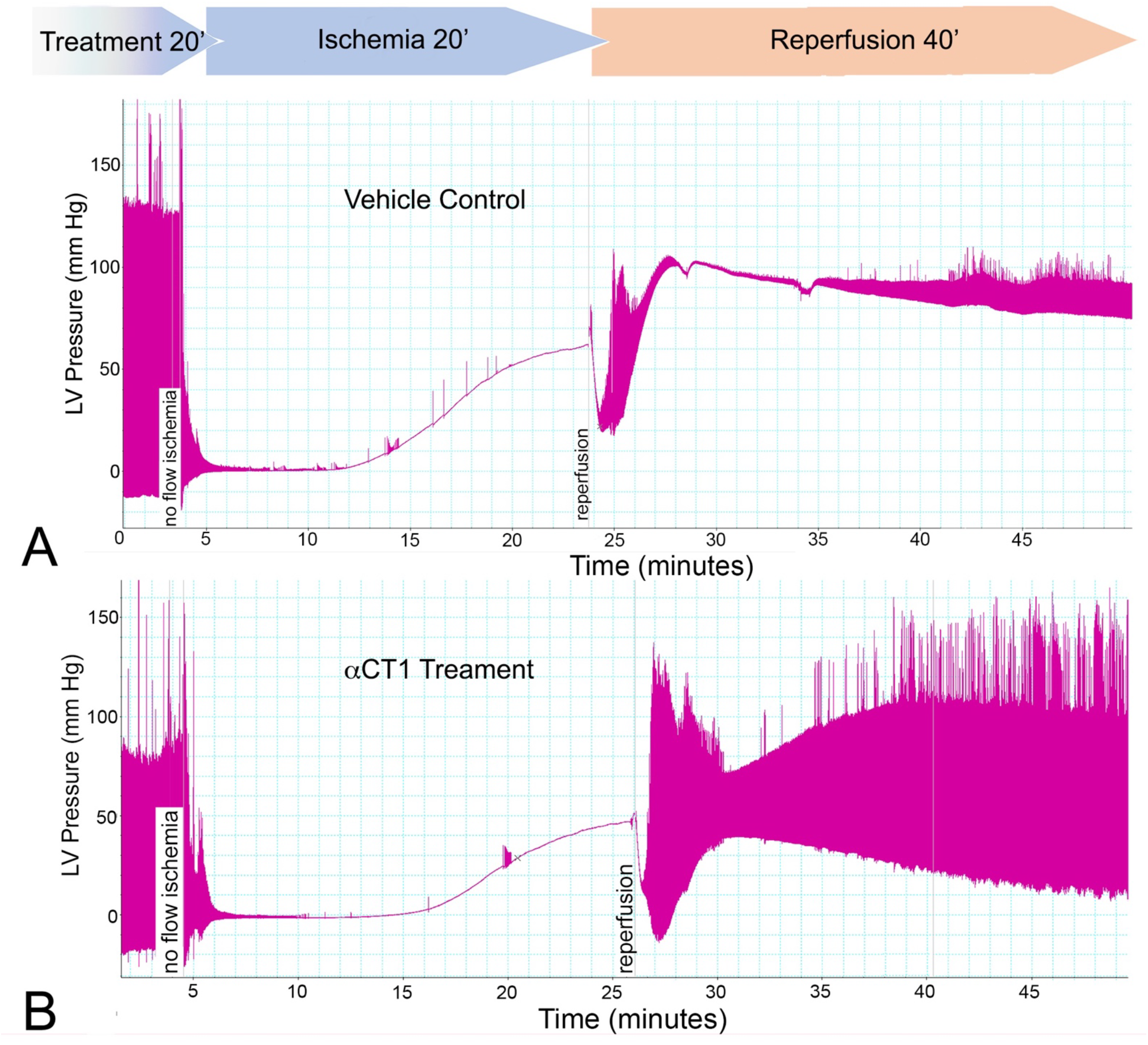
Pre-Ischemia treatment with peptides competent to interact with Cx43 CT protect hearts from ischemia-reperfusion (I/R) injury. Langendorff I/R protocols were performed on adult mouse hearts instrumented to monitor LV function (protocol in supplemental Fig. 1). Representative pressure traces from hearts from: (A) Vehicle control and (B) 10 μM αCT1 infused hearts. Note that the αCT1 treatment results in notable recovery of LV function during reperfusion.

The effects of the αCT1 and the αCT1-variants on left ventricular (LV) systolic and diastolic contractile function showed a striking correlation with the Cx43 CT ability of peptides (Fig. 7). Whereas the non-Cx43 CT interacting peptides M1 AALAI and M3 DDLAI showed no ability to improve recovery of either systolic (Figs. 7A-C) or diastolic (Figs. 7D-F) LV contractile performance during reperfusion, hearts pre-treated with the Cx43 CT-interacting peptides αCT1, αCT11 and αCT1-I demonstrated significant functional recovery after I/R injury, compared to vehicle control mice (Figs. 7A-G). Further, as αCT1-I is able to interact with Cx43 CT, but not PDZ2, the results suggested that ZO-1 binding was dispensable for induction of functional cardioprotection. Importantly, all Cx43 CT-binding peptides resulted in highly significant 3 to 5-fold improvements in functional recovery of LV contractile function during reperfusion following ischemic injury relative to vehicle control and the non-Cx43 CT interacting peptides (Fig. 7G). In line with the observations of the *in vitro* kinase assays (Fig. 3), LV samples taken for Western blotting following pre-ischemic treatment of Langendorff-perfused mouse hearts with αCT1, αCT11, αCT1-I showed significant increases in phosphorylation at the Cx43 PKCε-consensus locus S368 relative to vehicle control perfused hearts (Fig. 7H). By contrast, hearts exposed to peptides not competent to interact with Cx43 CT (i.e., M1 AALAI and M3 DDLAI), uniformly showed no propensity to upregulate S368 phosphorylation (Fig. 7H).

**Figure 7.**
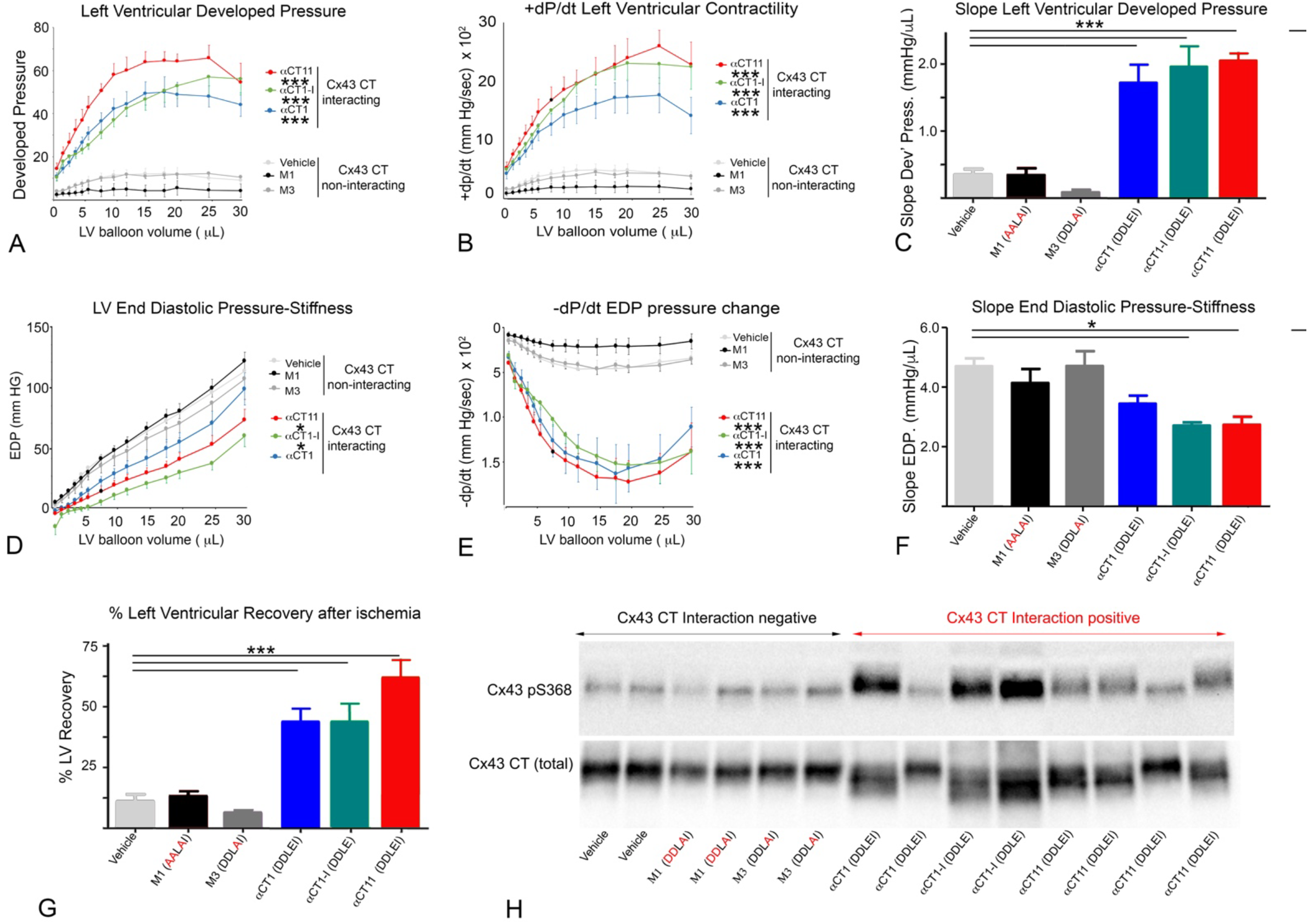
Pre-Ischemic treatment with peptides interacting with Cx43 CT protect hearts from ischemia-reperfusion injury in association with increased pS368 in LV myocardium. Langendorff ischemia-reperfusion (I/R) injury protocols were performed on adult mouse hearts instrumented to monitor LV contractility (protocol in supplemental Fig. 1). LV Systolic responses are shown in **7A-C**: **(A)** Plots of left ventricular (LV) systolic developed pressure against balloon volume**; (B)** LV maximal rate of tension development (+dP/dt) against balloon volume; **(C)** Maximal systolic elastance (E_max_) – i.e., the slope from (A)**; (D)** Plots of LV end diastolic pressure (EDP) against balloon volume; **(E)** Maximal rate of relaxation (-dP/dt) against balloon volume; **(F)** Stiffness, the reciprocal of the slope from (D); (**G)** Percentage of LV contractile function recovery post-ischemia relative to baseline level. Data shown are mean ± S.E. N=4-8. *p<0.05, ***p<0.001, N=4-8 hearts/group. **H)** Blots of Cx43-pS368 (top) and total Cx43 (bottom) of LV samples infused with peptide for 20 minutes according to the protocol in supplemental figure 1. For hearts used in Western blots, the protocol did not proceed to the ischemia and reperfusion phases, being terminated after the peptide infusion step. Only those peptides competent to interact with Cx43 CT increase pS368 levels relative to total Cx43 above vehicle control.

### Post-Ischemic Treatment with the 9mer peptide αCT11 preserves LV Function

αCT1 is in Phase III clinical testing in humans for pathologic skin wounds ^19^. The results of figure 7 indicated that pre-treatment with Cx43 CT binding peptides provided protection from injury in the ex vivo model studied. However, to be clinically useful to patients, such as those suffering a myocardial infarction, a drug would typically need to be given after an ischemic insult to the heart, i.e., after a myocardial infarction has been diagnosed. We thus treated hearts during the reperfusion phase following ischemic injury with αCT1, but determined that this did not result in significant recovery of LV function (data not shown). As αCT1 showed no evidence of post-infarction efficacy, we decided to explore an alternative approach. It was notable that the most striking recovery of post-ischemic LV function resulted from pre-ischemic treatment with the 9mer Cx43 CT-binding peptide αCT11 (Table 1). This is illustrated in Figure 7A, where the curve for LV developed pressure for αCT11 conspicuously overarches that of the other two Cx43-interacting peptides, αCT1 and αCT1-I. This can also be observed in Figure 7F, where the % of LV function recovery associated with αCT11 pre-infusion significantly exceeds that of αCT1 or αCT1-I (p<0.05). Based on these results suggestive of increased potency, we sought to determine whether αCT11 might have a post-ischemic cardioprotective effect.

αCT11 demonstrated an ability to significantly improve recovery of both systolic (Figs. 8A-C) and diastolic (Figs. 8D-F) LV contractile performance when infused in hearts during the reperfusion following ischemic injury. The level of cardioprotection achieved by this post-ischemic treatment was not as high as when αCT11 was provided prior to insult, but it was similar to that achieved for pre-ischemic treatment with αCT1.

**Figure 8.**
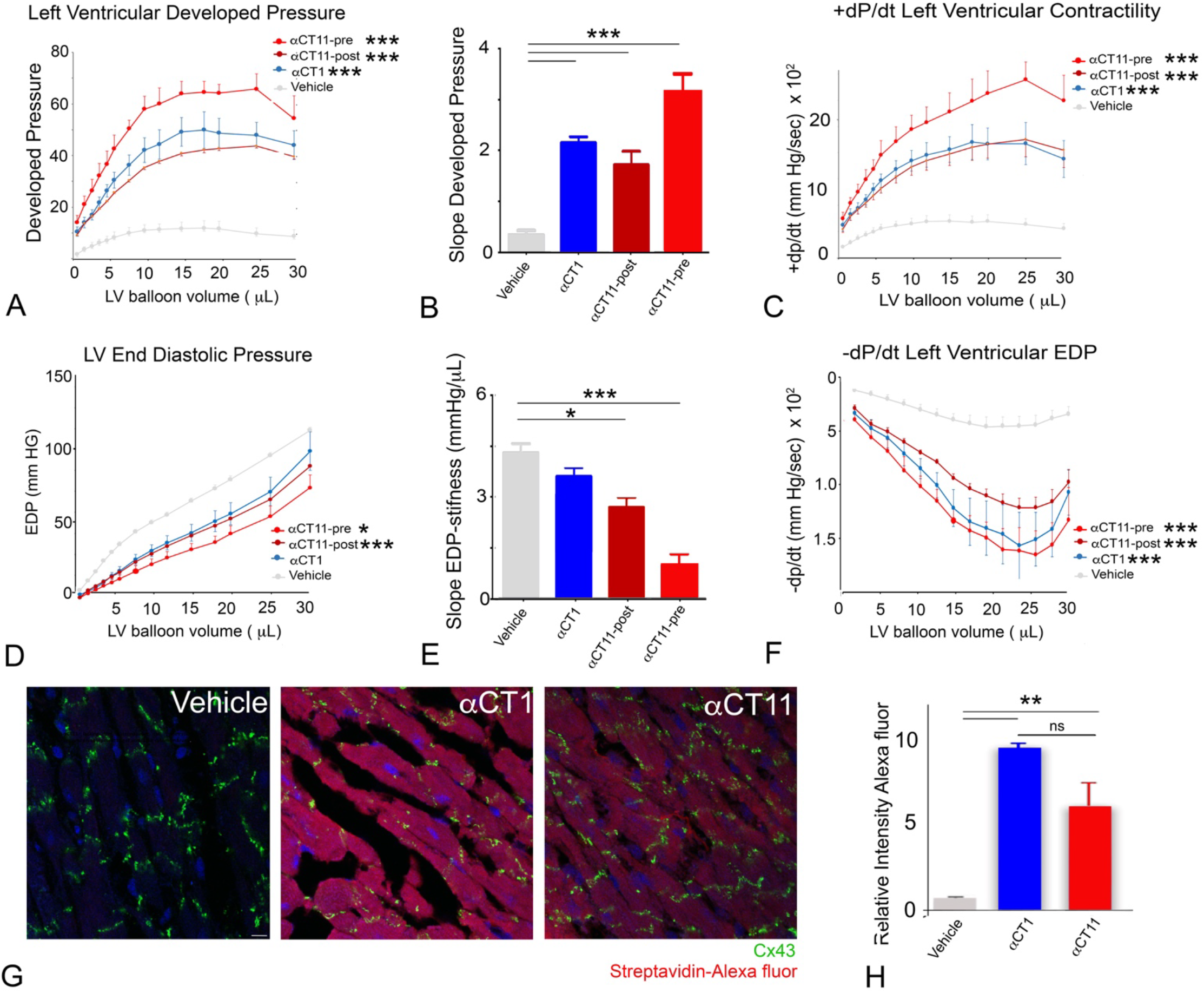
Pre- and Post-Ischemic treatment with αCT11 protect hearts from ischemia-reperfusion injury. Langendorff I/R protocols were performed on adult mouse hearts instrumented to monitor LV contractility. Protocol in supplemental Fig. 1, except that a 20-minute peptide infusion was begun after ischemic injury at the initiation of reperfusion. **(A)** Plots of left ventricular (LV) developed pressure against balloon volume; **(B)** Maximal systolic elastance (E_max_), the slope from (A); **(C)** Maximal rate of tension development (+dP/dt) against balloon volume; **(D)** Plots of end diastolic pressure (EDP) against balloon volume; **(E)** Stiffness, the reciprocal of the slope from (D); **(F)** Maximal rate of relaxation (-dP/dt) against balloon volume. * p<0.05, *** p<0.001, N=4-8. **G)** Laser scanning confocal microscopic fields from sections of Vehicle control, αCT1, and αCT11 group hearts stained for Cx43 (green), nuclei (DAPI-blue), and Alexa647-conjugated streptavidin (red). **H)** Average intensities of biotinylated peptide (indicated by streptavidin Alexa647 fluorescence intensity level relative to background) in Vehicle control, αCT1, and αCT11 groups. ** p<0.05; not significant (ns) N=5 hearts/group. Scale bar = 5 μm.

Given that αCT11 is missing a cell penetration sequence we were curious to determine whether the 9mer peptide (MW=1110 Daltons) was being taken up into cardiomyocytes. We thus examined uptake of αCT11 in ventricular muscle in mouse hearts that had been perfused with a biotinylated αCT11 under the protocol summarized in supplementary Figure 1. Cardiomyocytes showed robust uptake of the 9mer αCT11 sequence, as detected by fluor-conjugated streptavidin (Fig. 8G), and as compared to vehicle control perfused hearts. Relative to vehicle control, quantified levels of uptake of αCT11 in cells were comparable to those of αCT1, as indicated by measurement of relative fluorescence intensity levels in ventricular myocardial tissues (Fig. 8H).

## DISCUSSION

This study demonstrates that mimetic sequences incorporating the CT-most nine amino acids of Cx43 (amino acids R374 to I382) complex with the Cx43 CT – including the H2 sequence located between amino acids D340 and D360 of Cx43. This interaction appears to cause disruption of polypeptide structure, which in turn is associated with increases in a PKC-mediated phosphorylation in a serine residue at position 368 of Cx43 - S368. Moreover, evidence is provided that the cardioprotective properties of Cx43 CT mimetic peptides, such as αCT1, may be explained to a significant degree by their propensity to interact with the Cx43 CT. This conclusion is supported by results indicating that a Cx43 CT-binding competent peptides αCT1, αCT1–I and αCT11 preserve LV function following ischemic injury, whereas Cx43 interaction deficient variants of αCT1, M1 AALAI and M3 DDLAI, do not. Interestingly, αCT1 and H2 represent two spatially distinct sequences on the CT of native Cx43 molecules. Thus, the data point to the possibility of interactions between or within Cx43 molecules in vivo that may be involved in regulating Cx43 phosphorylation – potentially doing so by controlling accessibility of PKCε to its Cx43 CT substrate.

Our observations on the relationship between PKCε-mediated phosphorylation of S368 and cardioprotection are consistent with a longstanding literature ^6, 7, 11, 32–39^. Phosphorylation of Cx43 at S368 is correlated with reduced activity of Cx43-formed hemichannels ^6, 9, 10, 40^. Pro-inflammatory and injury spread signals resulting from unregulated opening of hemichannels in the myocyte sarcolemma are thought to be determinants of the severity of ischemia reperfusion damage to the heart ^41–52^. Cx43 activity and pS368 phosphorylation events associated with mitochondrial membranes have also been linked to I/R injury severity ^42, 53, 54^. Interestingly, it has been reported that Cx43 CT sequences incorporating the Cx43 H2-binding sequence of interest herein result from alternative translation of the GJA-1 gene (Smyth et al 2013, PMID: 24210816). These include a 20 kDA isoform, termed GJA1-20k, which has been found to be enriched at the interface between mitochondria and microtubules ^55^. Similar to the results achieved with synthetic Cx43 CT mimetic sequences here, exogenous provision of GJA1-20k reduces infarct size in mouse hearts subjected to I/R injury ^56^. Ongoing experiments would usefully address the extent to which treatment regimens based on αCT1-based peptides and GJA1-20k share aspects of molecular mechanism.

The pH-dependent gating of Cx43-formed channels has been proposed to involve the Cx43 CT in a ‘‘ball-and chain’’ mechanism ^57, 58^. The demonstration that the CT-most 10 amino acids of Cx43 (S373-I382 aka CT10) interacts with a region of the cytoplasmic loop domain of Cx43 referred to as L2, resulting in channel closure under acidic conditions, provides evidence supporting this hypothesis ^59^. We demonstrate in the current paper that a near-identical sequence to CT10 contained in αCT1 (i.e., R374-I382), also interacts with the H2 sequence of Cx43, doing so via precisely the same negatively charged amino acids required for L2 interaction ^20^. In addition to the shared affinity of the CT-most 9 amino acids of Cx43 for L2 and H2, comparison of L2 and H2 indicate other notable parallels. The L2 and H2 sequences of Cx43 have related secondary structures, both being marked by short stretches of α-helix. Further, L2 and H2 incorporate a pair of lysine (KK) residues. As we demonstrate herein, these lysines are essential for αCT1 interaction, as substitution of K345 and K346 with neutral glutamines, as in the Cx43 CT QQ/KK construct, results in a loss of αCT1 binding to H2.

Taken together, the evidence suggests that the nine amino acid CT sequence of Cx43 mimicked by αCT1 is a multivalent ligand that participates in a number of protein-protein interactions. In addition to affinity for L2 and H2, this short segment of Cx43 includes the PDZ-binding-ligand necessary for linkage to ZO-1 ^14, 60, 61^, as well as amino acids required for interaction with 14-3-3 theta ^62^. Immediately proximal are consensus recognition sites for AKT (S373) ^63^, PKCε (S368) ^64, 65^ and T-cell protein tyrosine phosphatase ^66^. The potential for complex patterns of protein-protein interaction occurring within such short stretch of the Cx43 primary sequence raises intriguing possibilities for future work. Of relevance are questions as to the timing of association of partnering proteins during the Cx43 life-cycle and the nature of the secondary and tertiary structures involved. With respect to the latter, it would be interesting to explore whether PKCε accessibility to S368 could be governed by intramolecular interactions within Cx43 or involves intermolecular interactions between Cx43 molecules. For example, dimers of Cx43 molecules involving overlapping interactions between the CT-most amino acids of one Cx43 molecule and the H2 domain of an adjacent Cx43 within a connexon, and changes in the avidity of this dimerization, suggests itself as one hypothesis as to how PKCε accessibility to S368 might be regulated.

Our results indicate that the Cx43 CT-binding activity of αCT1, and not ZO-1 PDZ2 interaction, explains the cardioprotective effects of αCT1, at least in the model studied here. Whilst Cx43-ZO-1 interaction does not appear to have been a direct factor in the ex vivo model studied, potential roles for ZO-1 in regulating Cx43 phospho-status and hemichannel availability in vivo, including during ischemic injury, should not be discounted. ZO-1 is located at the edge of Cx43 GJs in a specialized zone of cell membrane known as the perinexus ^63, 67, 68^. In earlier studies, we have shown that high densities of hemichannels are found in this peri-junctional region ^69, 70^ and that PDZ-based interactions between ZO-1 and Cx43 govern the rate at which undocked connexons dock with connexons from apposed cells to form gap junctional channels, thereby regulating GJ size, as well as hemichannel availability within the cell membrane ^13, 14^. Recent work by two other groups have provided data supporting this hypothesis, and have also shown that phosphorylations at Cx43 S368 and S373 are central to how ZO-1 controls the accrual of perinexal hemichannels to the GJ ^63, 71^. The potential for regulatory interplay between PKC-ε and ZO-1 at the Cx43-CT is further suggested by earlier studies indicating that the presence of ZO-1 PDZ2 domain in the test tube-based PKC assay efficiently acts as a competitive inhibitor of αCT1 enhancement of S368 phosphorylation ^11^.

A key question raised by our study is whether the αCT1 Cx43-targeting mechanism determined as necessary for preservation of LV function also explains the primary mode-of-action of this therapeutic peptide in other tissues. In skin wounding experiments in mice and pigs, αCT1 has been shown to reduce inflammation, increase wound healing rates and decrease granulation tissue formation ^12, 24^. In related observations in Phase II clinical testing of humans, αCT1 treatment increased the healing rate of slow-healing skin wounds, including diabetic foot ulcers and venous leg ulcers ^16, 18^. Given the current results in heart, it will be of interest to determine whether the mode-of-action of αCT1 in wounded skin also involves Cx43 CT interaction and/or increased pS368. As the GAIT1 Phase III clinical trial on αCT1 moves forward on more than 500 patients with diabetic foot ulcers ^19^, such insight on molecular mode-of-action will be useful in understanding the basis of any clinical efficacy identified in humans, as well as a step in building a safety profile for this therapeutic peptide.

Of further clinical translational note are our findings on the cardioprotective effect of post-ischemic treatment by the short αCT1 variant αCT11 - a result that may have clinical implications. Interestingly, αCT11 does not have a cell penetration sequence, but it nonetheless appears to be internalized by LV cardiomyocytes after intravascular perfusion in the ex vivo model used herein. The mechanism of this cellular uptake is presently under study by our group, but it may be explained by the small size (MW=1110 Daltons) and linear, random coiled-coil 3D structure of αCT11 – see Figure 2A. Neijssen and co-workers reported that linear peptides with molecular masses below 1800 Daltons readily diffuse through Cx43-formed channels ^72^. Given αCT11 has a molecular mass well below 1800 Daltons, and that hemichannel opening is induced by ischemic insult ^41^, the interesting prospect is raised that αCT11 does not have far to go to reach its target (i.e., the cytoplasmic CT domain of Cx43), potentially doing so by simply transiting open Cx43 hemichannel pores in the myocyte sarcolemma. Future work would usefully test this hypothesis, as well as undertake further testing of αCT11 in preclinical models of cardiac I/R injury *in vivo* as a prelude to Phase I testing of this novel therapeutic peptide in human patients with acute myocardial infarction.

## ACKNOWLEDGMENTS

We thank Greg Hoeker and Randy Strauss for reading and providing suggestions on the manuscript. We thank Linda Collins for her editorial contribution to this manuscript.

## Contributions

JJ, JAP, HH, GB, KS, SP, DH, FM and RG were responsible for experimental design and data interpretation.

JJ, JAP, HH, JI, JJ, and ZW conducted experiments.

RG was largely responsible for manuscript preparation, with editing assistance from JJ, HH, GB, DH, ZZ, KS, SP, FM.

---

**Supplemental Figure 1.**
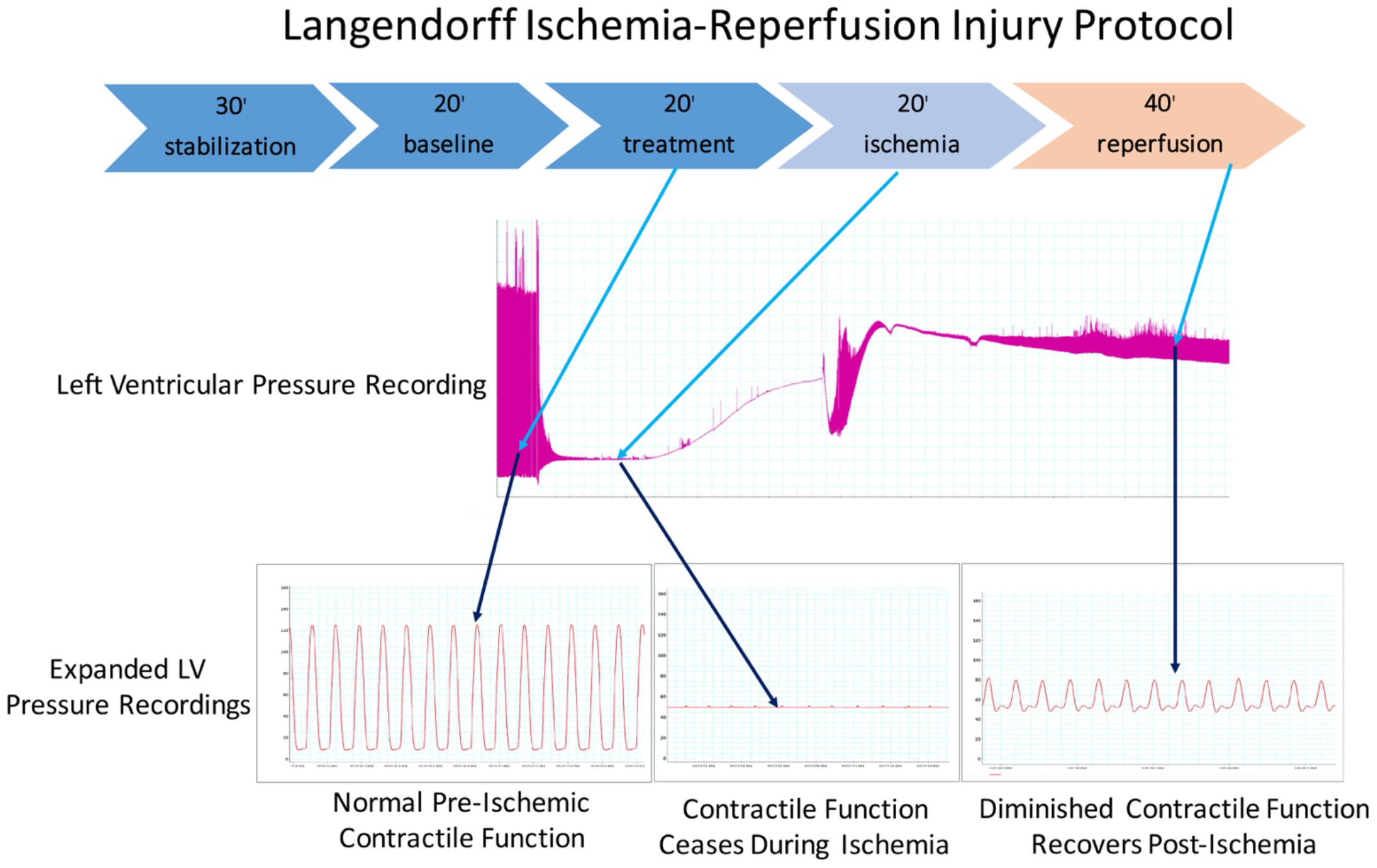
Ischemia reperfusion injury model/protocol. The protocol involved a 20-minute period of no flow ischemia period followed by 40 minutes of reperfusion, LV contractile function was monitored throughout the whole process. For treatment, peptides were infused into hearts over a 20-minute period just prior to the ischemic episode. Expanded representative pressure traces for each of these phases are shown below.

**Supplemental Figure 2.**
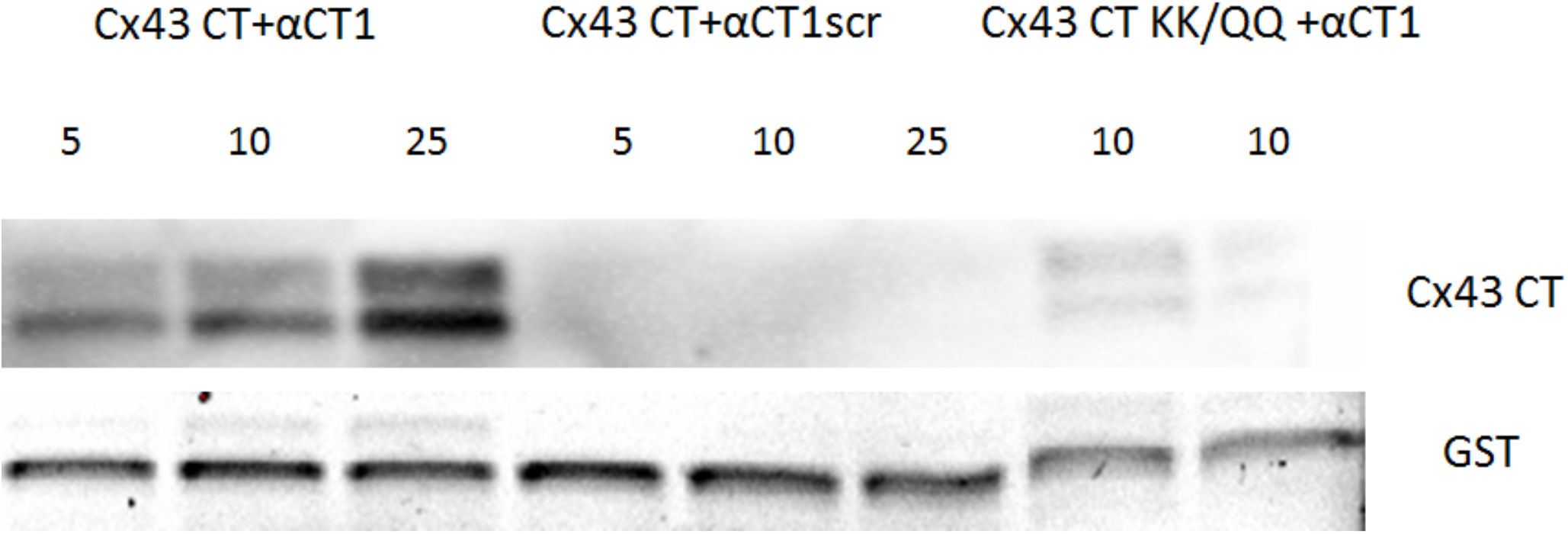
Blots of EDC cross-linked products of kinase reaction mixtures containing GST-Cx43 CT, GST-Cx43 CT QQ/KK in which the lysine (K) residues were mutated to neutral glutamines (Q), PKC-ε and αCT1 (at 5, 10 and 25 μM) and a scrambled αCT1 (M4 scr) variant at the same concentrations. Only αCT1 is seen to be covalently linked by EDC to Cx43 CT in a concentration-dependent manner.

**Supplemental Figure 3.**
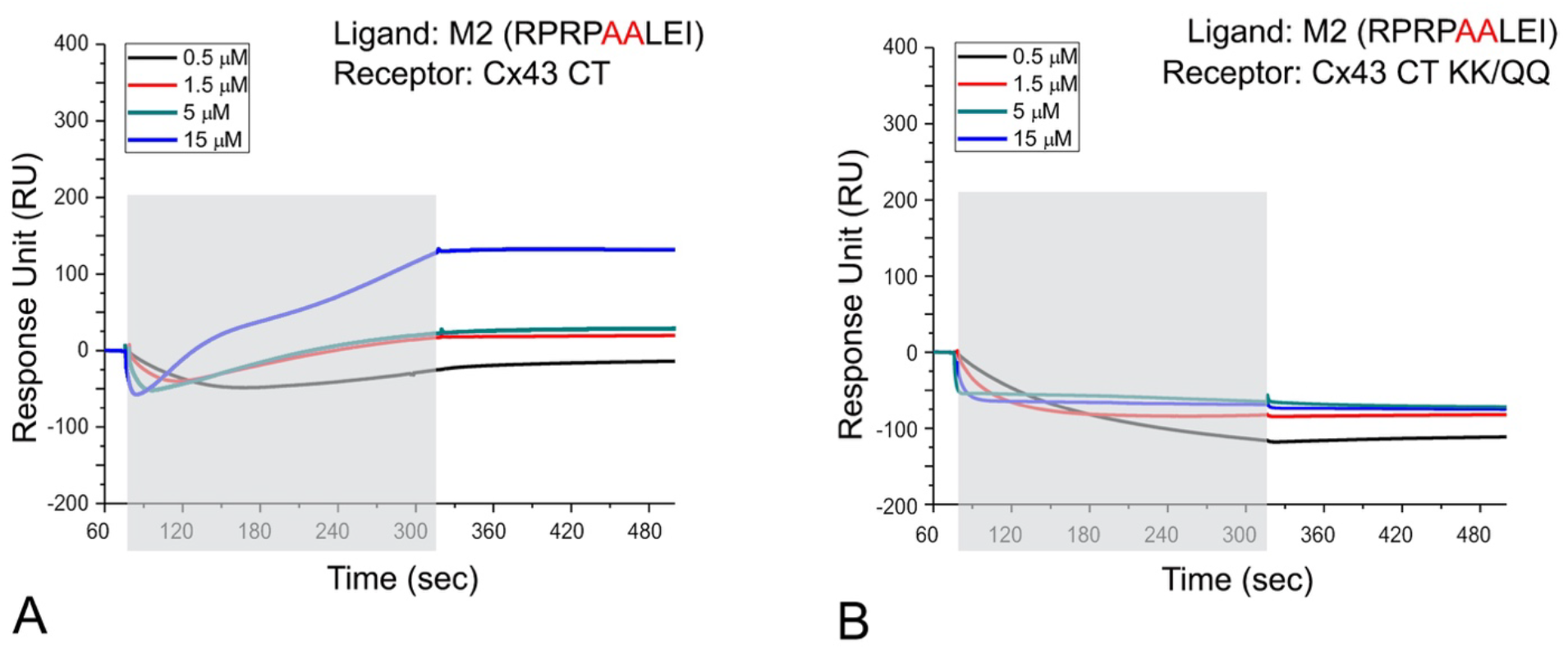
The αCT1 variant peptide M2 AALEI shows limited ability to bind Cx43 CT. SPR was used to analyze interactions of biotin-M2 AALEI with the Cx43 CT (**A**) and Cx43 CT-KK/QQ (**B**) as respective analytes. The mean of three runs is plotted for each analyte concentration. The exposure of the sensor chip to the specific analyte is indicated by the gray area.

